# Speaq 2.0: A Complete Workflow for High-Throughput 1D NMR Spectra Processing And Quantification

**DOI:** 10.1101/138503

**Authors:** Charlie Beirnaert, Pieter Meysman, Trung Nghia Vu, Nina Hermans, Sandra Apers, Luc Pieters, Adrian Covaci, Kris Laukens

**Author notes:** We dedicate this paper to our colleague, Prof Sandra Apers, who passed away much too early on February 5th, 2017.

## Abstract

Nuclear Magnetic Resonance (NMR) spectroscopy is, together with liquid chromatography-mass spectrometry (LC-MS), the most established platform to perform metabolomics. In contrast to LC-MS however, NMR data is predominantly being processed with commercial software. This has the effect that its data processing remains tedious and dependent on user interventions. As a follow-up to speaq, a previously released workflow for NMR spectral alignment and quantitation, we present speaq 2.0. This completely revised framework to automatically analyze 1D NMR spectra uses wavelets to efficiently summarize the raw spectra with minimal information loss or user interaction. The tool offers a fast and easy workflow that starts with the common approach of peak-picking, followed by grouping. This yields a matrix consisting of features, samples and peak values that can be conveniently processed either by using included multivariate statistical functions or by using many other recently developed methods for NMR data analysis. speaq 2.0 facilitates robust and high-throughput metabolomics based on 1D NMR but is also compatible with other NMR frameworks or complementary LC-MS workflows. The methods are benchmarked using two publicly available datasets. speaq 2.0 is distributed through the existing speaq R package to provide a complete solution for NMR data processing. The package and the code for the presented case studies are freely available on CRAN (https://cran.r-project.org/package=speaq) and GitHub (https://github.com/beirnaert/speaq).

**Author summary:** We present speaq 2.0: a user friendly workflow for processing NMR spectra quickly and easily. By limiting the need for user interaction and allowing the construction of workflows by combining R functions, metabolomics data analysis becomes fully reproducible and shareable. Such advances are critical for the future of the metabolomics field as it needs to move towards a fully open-science approach. This is no trivial goal as many researchers are still using black-box commercial software that often requires manually doing several steps, thus hampering reproducibility. To encourage the shift towards open source, we deliberately made our method usable for anyone with the most basic of R experience, something that is easily acquired. speaq 2.0 allows a stand-alone analysis from spectra to statistical analysis. In addition, the package can be combined with existing tools to improve performance, as it provides a superior peak picking method compared to the standard binning approach.

## Introduction

1D NMR spectroscopy has been a popular platform since the early days of metabolomics. Although less sensitive than the complimentary and more common LC-MS technology, NMR it has its merits. For one, it is an unparalleled technique in determining the structure of unknown metabolic compounds. Furthermore, because it is a non-destructive technique, samples can be reanalyzed later or can be used in a different spectroscopic analysis such as mass spectrometry. Also, an NMR spectroscopy experiment requires little sample preparation compared to LC-MS, thus limited unwanted extra variability is introduced. Finally, the results of an NMR analysis are less dependent on the operator and instrument used. All these factors make 1D NMR spectroscopy a technique with a relatively high reproducibility and rather minimal researcher bias [1]. There are however also drawbacks associated with this technique. First, the aforementioned low sensitivity is an important issue as the dynamic range in real biological samples surpasses the NMR detection range. This is particularly problematic when the goal is to identify an unknown metabolite with a low concentration.

To get the best of both worlds, combining large scale LC-MS analysis with NMR spectroscopy has been presented as an option to yield valuable novel insights in metabolic pathways and biomarkers [2–4]. From a data processing perspective, this combination is not trivial. The data analysis of NMR is not as automated as LC-MS data analysis, which can rely on open-source solutions like XCMS [5]. Most NMR data analysis is still performed on commercial software [6]. While the reproducibility of the NMR experimental techniques is high, the data analysis still requires a substantial degree of user intervention. This results in the possible introduction of bias and lower research reproducibility, meaning that the data analysis can not be easily replicated by others. The absence of standardized and automated NMR metabolomics workflows is the main culprit despite recent progress. This progress includes the proliferation of R packages for NMR analysis like speaq [7], BATMAN [8], muma-R [9], ChemoSpec [10], and specmine [11]. But also other widely used frameworks are recent developments, such as MetaboAnalyst [12–14] a web server for LC-MS and NMR metabolomics analysis, Bayesil [15], another web based approach and MVAPACK [16], a GNU Octave toolbox to process NMR data starting from the raw FID signal. Not all these NMR analysis tools are applicable to all research setups. Some serve only specific purposes like BATMAN [8] for example which is aimed at obtaining concentration estimates for known metabolites from the raw spectra. However, a lot of untargeted experiments are in search for not only known metabolites, but also unknown ones. These experiments require tools that can process large amounts of data in a scalable way.

A typical workflow for NMR spectral analysis consists of several pre-processing steps, such as baseline correction, raw spectra alignment, spectra summarization and grouping. This is then followed by statistical analysis. The spectra summarization step and the alignment/grouping step are most time consuming. In the spectra summarization, all the experimental measurement points are transformed into a small number of features, which are more suited for automated analysis. Multiple spectra summarization techniques exist, each with their own advantages and drawbacks [18]. The specific method that is chosen can result in user-introduced bias and low reproducibility. This is the case for the most common used summarization approach: the so-called binning or bucketing method [19]. This method was introduced to compensate for small spectral shifts between samples. It allows to vastly reduce the number of measurements points to a limited number of variables (the bins) in one single, automated step [20]. There are however major drawbacks to this method that have a profound influence on the results [21]. In particular, it is not straightforward to define the boundaries of the bins in crowded spectra. Automating this process may lead to splitting up small but relevant peaks. Manually checking the bins on the other hand is extremely tedious and manually changing the boundaries can forfeit any attempt for reproducibility. Several methods have been proposed to tackle the bin boundary issue [22–24], but this is not the only concern. Loss of information is intrinsically linked to the binning approach as entire bins are simply summed together.

At the end of an analysis based on the binning approach, when several bins are found to be interesting, it is still necessary to revert to the raw spectra to manually check the intervals. This is necessary to find the ppm values of the actual peaks of interest that can then be used to query a database, like HMDB [17]. This introduces yet another point where user intervention is required, which slows down the whole process and hampers the use of an automated workflow.

In this paper we present the speaq 2.0 method. The underlying core paradigm is to efficiently summarize spectra with little user interaction, high speed and most importantly little loss of information whilst greatly reducing the dimensions of the data. The overall aim however, is not to construct yet another all-encompassing package for NMR analysis, but more importantly, to construct a method that can help established tools for NMR data analysis, like MetaboAnalyst [14], by improving performance, analysis quality and reproducibility. This is achieved by improving the quality of the peak lists which are the starting points for MetaboAnalyst [14] or muma-R [9]. By automating the important peak picking step in the NMR analysis workflow, less researcher bias is introduced hereby greatly improving reproducibility. The automation potential of the package makes it suitable for the fast analysis of NMR spectra in a way that is very comparable to how LC-MS spectra are analyzed. In the future, this method will effectively be used for high-throughput analyses in which LC-MS and NMR data are combined to achieve better results. Nonetheless, a complete standalone analysis pipeline is incorporated with the focus on user-friendliness. This is to allow also non-expert users to be able to work with open-source tools instead of the black-box proprietary software.

The basic proposition of speaq 2.0 is to use peaks instead of raw spectra. The peak-picking process is achieved with wavelets. Specifically the Ricker wavelet, also called the Mexican hat wavelet, is used to mathematically represent the peaks in the spectra in such a way that a large reduction of variables is achieved with very little loss of information. Only a few values capture the peak information that is contained in the tens or hundreds of raw data points describing the peak in the original spectra. Besides the data reduction the additional advantage of using wavelets, and specifically the continuous wavelet transform, is that the need for baseline correction and smoothing is eliminated with no loss of sensitivity or increase in false positives [25, 26].

## Materials and methods

### Benchmark Data

To validate the presented approach two previously published benchmark datasets are reanalyzed:

1. The wine dataset by Larsen et al. [27] where 40 table wines (red, white & ros´e) are measured with 1H NMR spectroscopy. The focus of Larsen et al. was not to investigate differences between wines of different colour and origin, but merely to evaluate how pre-processing methods like alignment and interval selection can aid in chemometrics and quantitative NMR analysis [27]. Wine is a good example for evaluating alignment algorithms because of the often substantial differences in pH, which can cause large shifts in the NMR spectra. Because of this property, the wine dataset has been used to evaluate the performance of several alignment algorithms, like COW [27], icoshift [28] and CluPA [7].
2. The onion intake in mice dataset which originates from a nutri-metabolomics study by Winning et al. [29] in search of biomarkers for onion intake. 32 rats were divided into 4 categories each receiving a specific onion diet: control (0% onion), 3% onion residue, 7% onion extract and 10% onion by-product. Urine samples were collected during 24 hours and analyzed with proton NMR spectroscopy to characterize the metabolome of the different onion fed mice. More details can be found in [29].

Both datasets were originally made available by the University of Copenhagen at http://www.models.life.ku.dk/.

### Workflow

The NMR data analysis workflow of speaq 2.0 is depicted in Fig 1. Spectra serve as input, then peak picking with wavelets is applied to transform the spectra to peak data. These peaks are then grouped into features with the grouping algorithm. The resulting matrix of features and samples are then used in statistical analysis. The following section describes the individual steps in more detail.

**Fig 1.**
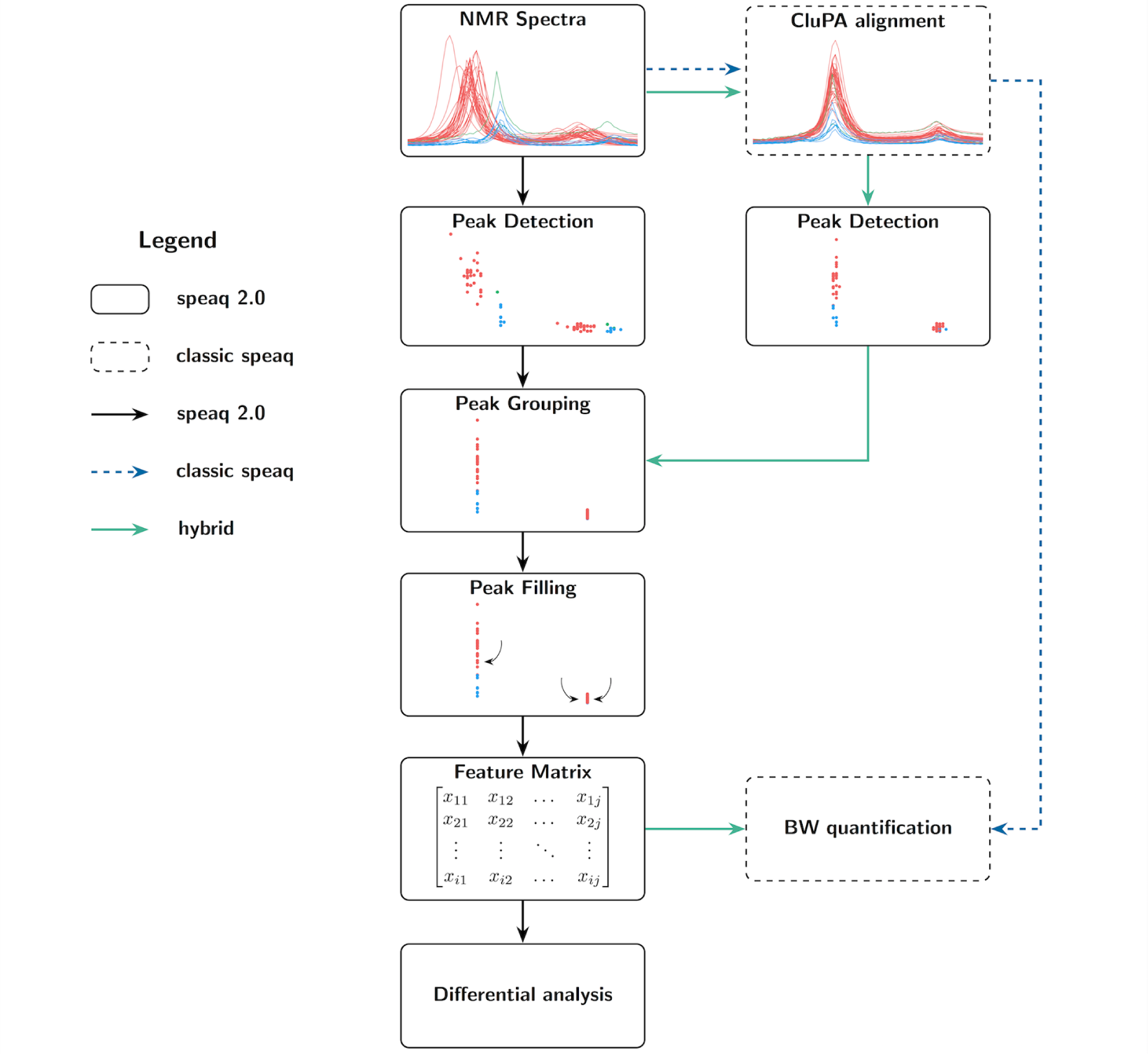
Possible workflows of speaq 2.0. The newly presented methods are standalone (black arrows) or can be used together with the CluPA alignment algorithm [7] (green arrows). It is still possible to perform an analysis based on raw spectra alone, as per the classic speaq analysis. In the new methods raw data is converted to peaks, and every peak is summarized with ppm location and width, intensity and SNR. These peaks are subsequently grouped and optionally peak filled (missed peaks in samples are specifically searched for). The resulting data is converted to a feature matrix that contains intensities for each peak and sample combination. This matrix can then be used in statistical analysis with built-in or external methods.

### Pre-processing steps

The input to the workflow consists of spectra in the intensity (y-axis) vs ppm (x-axis) format. This means that the free induction decay (FID) signal coming from the NMR spectrometer has to be converted to spectra by using the Fourier transform. In addition, before peak picking, the spectra can be aligned with the included CluPA algorithm [7] (the core of the first speaq version). Note that it is also possible to analyse spectra that have already been aligned with other methods like icoshift [28]. However, depending on the algorithm used, aligning raw spectra can result in the distortion of small peaks [30].

### Peak Detection: from spectra to features via wavelets

The Mexican hat or Ricker wavelet is used to perform the peak detection. It is a suitable wavelet because it resembles a peak by being symmetrical and containing only 1 positive maximum [25]. This peak detection method is inspired by the CluPA alignment algorithm in the original speaq software [7] where wavelets are used to find landmark peaks to aid in the alignment. The interaction with the wavelets relies on the MassSpecWavelet R package which performs the actual peak detection in the two-dimensional (position and scale) wavelet transform space by using ridge detection as per the method outlined by Du et al [25]. The result is a peak detection that is both sensitive to low and high intensity peaks and insensitive to background noise. After the peak detection, the spectra (intensity vs ppm data) are converted to a dataset with peakIndex and peakValue values. There is a direct link between a) ppm and peakIndex and b) intensity and peakValue whereby the peakValue is related to the amplitude of the wavelet that describes the peak. Note that this peakValue vs peakIndex dataset is substantially smaller than the original data.

### Peak Grouping

The peaks resulting from the wavelet peak detection are not perfectly aligned since no two peaks are exactly the same and different spectra can be shifted relative to each other. These shifts can be caused by differences in sample environment (pH, solvent, etc.) or differences in experimental conditions (temperature, magnetic field homogeneity). However, the aim is to go towards a feature dataset whereby a feature is defined as a group of peaks with at most one peak per sample belonging to that feature. This means the peaks have to be grouped with a single index value describing the group center. To group the NMR peaks we can make optimal use of the results of the wavelet based peak detection step. Not only ppm values but also signal-to-noise ratio and sample values can provide additional information to aid in the grouping. The hierarchical clustering based algorithm developed for grouping divides the samples in groups based on the distance matrix. It is illustrated with pseudocode in Fig 2 and a more detailed description can be found in the supplementary files.

**Fig 2.**
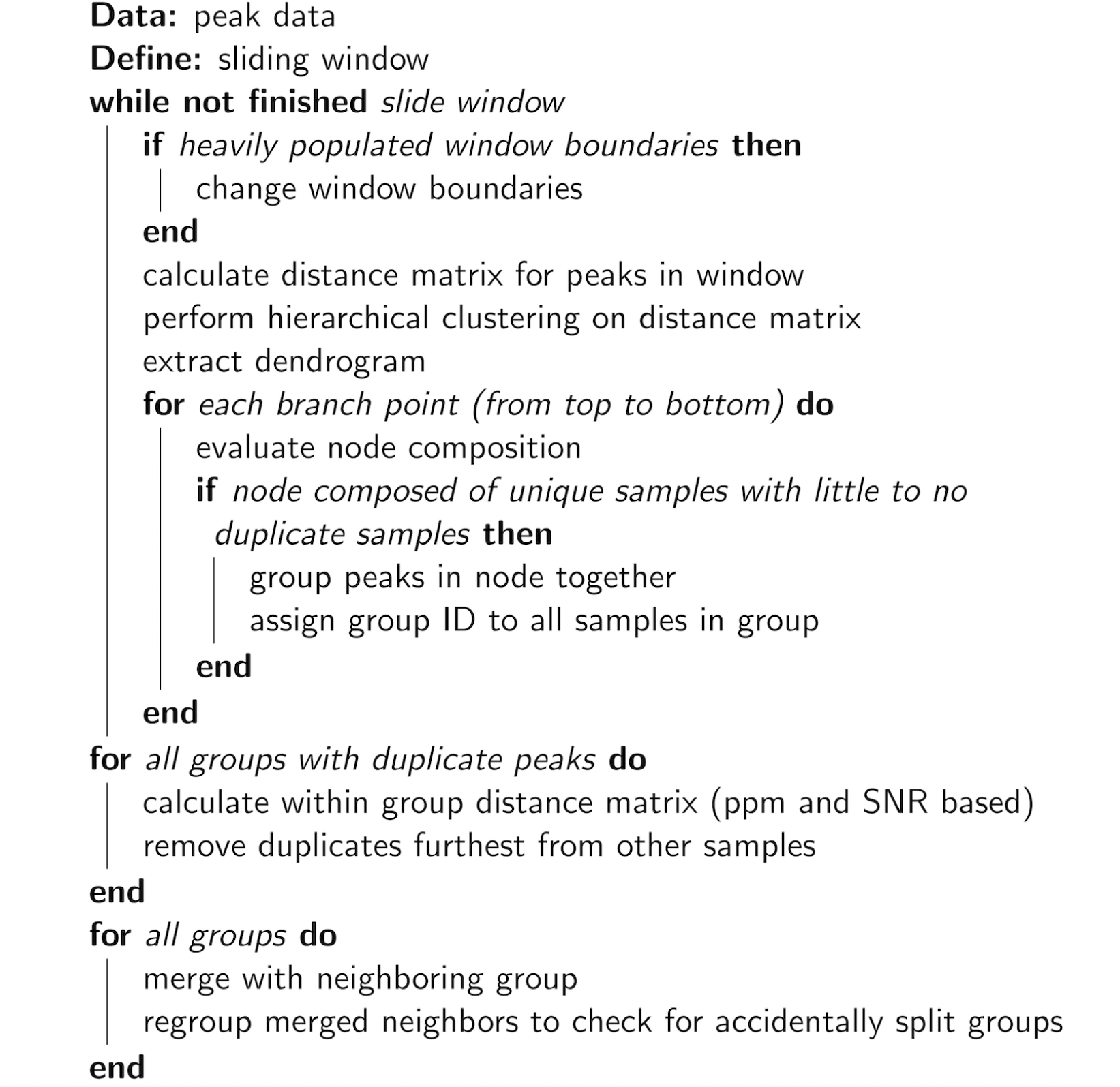
Grouping algorithm pseudocode.

Note that this method is designed to process data that is sufficiently well aligned. If this is not the case the method will most likely underperform because of the larger overlap between peaks. Nonetheless the method even works on data with non-trivial shifts between samples as is the case in the wine dataset [28]. Extremely shifted spectra can be aligned with existing methods such as CluPA [7], prior to peak detection.

### Peak Filling

The purpose of peak filling is to detect peaks that may have been missed in the first round of peak detection or that were deleted in one of the grouping steps. Because of the way the grouping is implemented it is advised to perform the peak filling step. The motivation for peak filling is that, when certain samples are represented in a peak group and other samples not, then it is not trivial to know if the signals are indeed absent or if some peaks where missed by the wavelets or deleted in the grouping step. If the peak is very large and prominent it can be assumed that the peak is simply not present in the missing sample. However, when the peak is small or close to a specified signal-to-noise ratio threshold then some peaks might end up missing while in fact they are present in the data. With peak filling, the information of the samples that are present in a feature is used to specifically search the raw data for peaks of missing samples. This results in more peaks, which in turn results in a more robust statistical analysis afterwards as less missing values have to be imputed.

### Statistical analysis

Following peak filling, the data can now be represented in the form of a matrix with samples for rows, features (peak groups) for columns and peak values in each matrix cell. Each of these peak values corresponds to the intensities of the original peaks as quantified by the wavelets. A huge number of techniques for univariate and multivariate statistics (e.g. PCA, PLS-DA) and machine learning (e.g. SVM, random forest) can be applied to this data matrix. A few methods have been directly integrated into the speaq 2.0 framework: a differential analysis method and a tool to perform scaling, transformations and imputations.

### Scaling and imputation

Before statistical analysis methods like PCA can be used the missing values in the data have to be imputed. This step is often done in tandem with the desired scaling method since otherwise data can artificially be created. For example, imputing zeros followed by z-scaling is not the same as z-scaling followed by imputing zeros. The last actually corresponds with imputing mean values. For both benchmark datasets zeros are used for imputing missing peak values in the data matrices as this indicates intensity 0, i.e. a non present peak. After imputing the scaling step is executed. Several methods for scaling are available in our framework, one of which is Pareto scaling [32]. This method is most suited for metabolomics since it reduces the effects large signals while keeping the data structure roughly intact. Pareto scaling is governed by the formula in Eq (1) with x_*j*_ the *j*^*th*^ feature vector containing the peak values *x*_*i,j*_ of all samples 1…*i*…*N* and *σ*_*j*_ the standard deviation of x_*j*_.

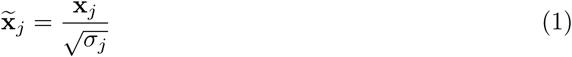

### Principal component analysis

Principal component analysis, or PCA, is an unsupervised multivariate analysis technique based on projections of the data onto new variables or dimensions called principle components. These principal components are orthogonal to one another and are combinations of the old variables in such a way as to maximize the variance (information) within a principal component. Plotting the data on these principal components, the so-called score plot, is often the first step in an analysis of large datasets as it can indicate groups, trends, or outliers. In addition, it was used in the original study of mice onion intake by Winning et al. [29] and therefore PCA is also used in our analysis to compare results.

### Differential analysis

A method to perform a differential analysis based on linear models is available in speaq 2.0. This function provides a way of identifying significant features with (adjusted) p-values. Specifically, for each feature 1,…, *j*,…, *K* consisting of peak values *y*_i,j_ of samples 1…*i*…*N* a linear model of the form

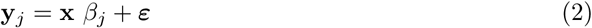

is constructed with x the response vector (N elements), *y*_*j*_ the *j*_*th*_ feature vector and *ε* the vector of errors *ε*_*i*_. Now for each *β*_*j*_ we can test whether there is a significant relationship between feature *y*_*j*_ and x by testing the hypothesis that *β*_*j*_ = 0 (two-tailed t-test). The *K* p-values can be used to find peaks significantly associated with the outcome vector. Several multiple testing corrections are available within the speaq 2.0 framework. For the purposes of this study, the stringent Bonferroni correction (default is Benjamini-Hochberg) was applied to all reported p-values. Note that in the case of only two classes this method is equivalent to the t-test.

### Metabolite identification

After statistical analysis the relevant peaks can be matched with the molecules responsible for these peaks. Several databases are available to facilitate the metabolite lookup, such as The Human Metabolome Database (HMDB) [17]. To obtain the metabolites for the onion intake in mice data the latest version of HMDB (3.6 [33]) was used. It is however not optimal to submit all significant peaks in a single query to this database since these peaks can come from different metabolites. HMDB works by matching the queried peaks to the database and then sorting the matched molecules according to their Jaccard index. For two sets the Jaccard index is the number of common elements (the intersection), divided by all the elements (the union), or in this specific case the number of matched peaks divided by all peaks in the query. Adding additional peaks from molecule B when trying to identify molecule A will dilute the results. To eliminate the problem of submitting peaks from multiple compounds, a correlation analysis is performed [34]. After all, in NMR spectra we can assume that peaks originating from the same molecule exhibit a similar behavior over all samples. Therefore the peaks that correlate strongly with each other are most likely to come from the same metabolite and can as such be submitted to HMDB simultaneously.

## Results

### Wine data

The first validation dataset concerns the NMR spectra of table wines. This is an often used dataset to evaluate alignment performance [7, 28]. The right half of Fig 3 illustrates the results from the peak detection and grouping procedures. Peak detection is applied to the raw spectra to convert the large raw measurement data matrix of 40 samples by 8712 measurements to a smaller matrix of 6768 peaks by 6 columns consisting of values describing the peaks. The data reduction after this step does not seem overwhelmingly large. However, it is important to realize that this is only a reduction in redundant information which is accompanied with almost no loss of information thanks to the wavelets. Furthermore, most of the correlation between consecutive measurement points in the spectral data is removed. After this step, the peaks are grouped, resulting in the same dataset as the peak data, but now each peak has been assigned to a group. Such a group consists of a collection of peaks with at most one peak per sample. The grouping method can theoretically under-perform when spectra are severely misaligned. However, for this dataset the grouping algorithm performs as expected, despite the severely shifted spectra. This grouped peak data can now be represented as a matrix, with groups as columns, samples as rows, and peak intensities as the matrix elements. The true data reduction becomes apparent now as there are only 207 peak groups, which correspond to the features used in further analysis. The original matrix of 40 by 8712 is thus converted to a matrix of 40 by 207.

**Fig 3.**
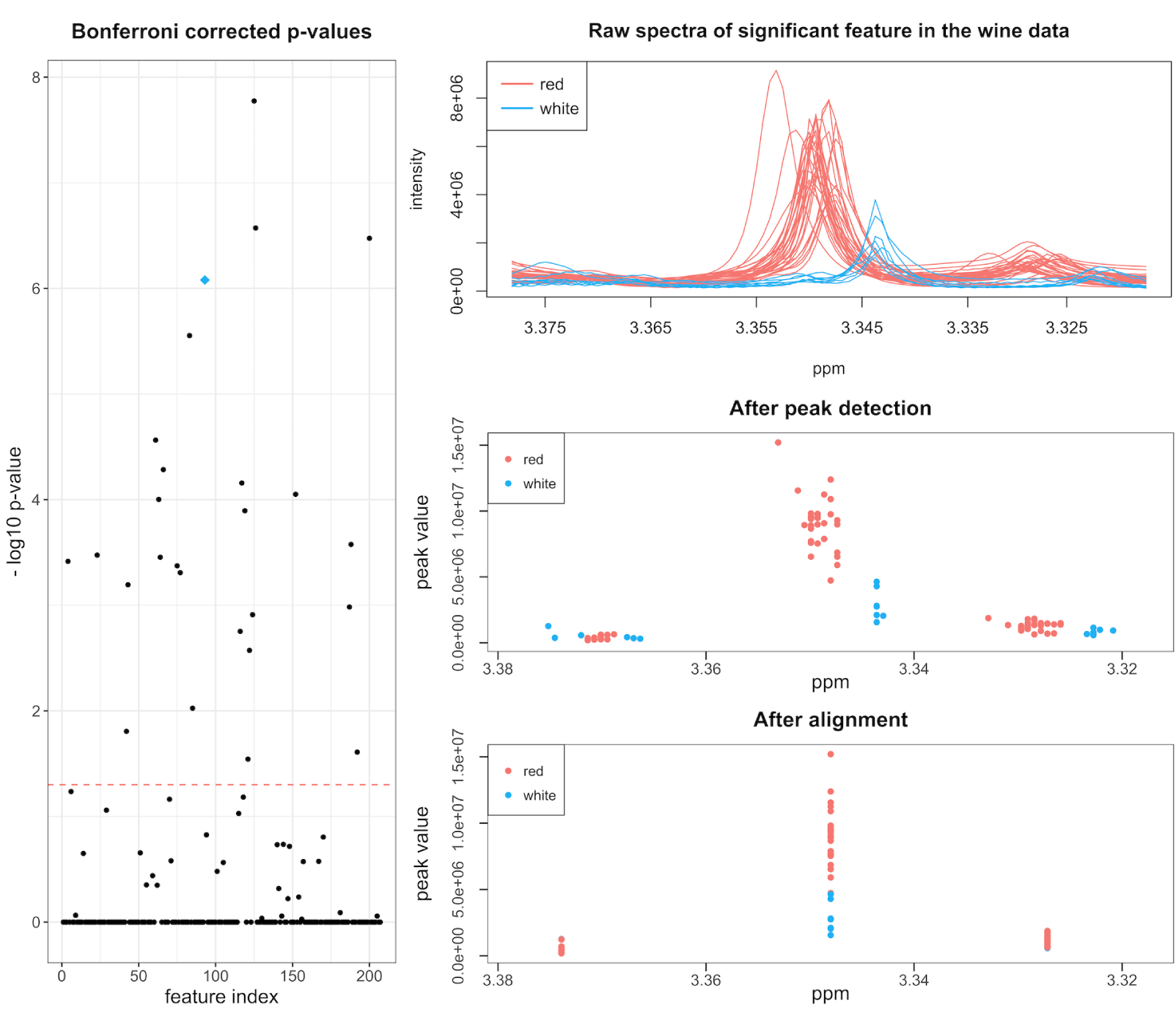
Visualization of Bonferroni corrected p-values. Numerous features have a corrected p-value below the significance threshold of 0.05 indicating that there is a significant difference between red and white wine. An example of a significant feature (indicated with the blue diamond) is represented on the right with its raw spectra (top), the data after peak detection (middle) and the data after grouping (bottom).

Next, we can identify those features that are associated with wine type. Before any analysis the data matrix is Pareto scaled and centered. The first step in a multivariate analysis is often principal component analysis (PCA). The results show that there is a clear difference between on one side red and on the other white and ros´e wines (see supplementary files). However, a differential analysis method incorporated into speaq 2.0 is more useful to identify the specific features that are different between the wine classes. Since a differential analysis is between two groups, the two samples that are neither red or white are excluded. The results of the differential analysis is a series of p-values, one for every feature, which indicate how useful each feature is in building a linear model that can discriminate between the two wine classes. These p-values are Bonferroni corrected to minimize the effects of multiple testing. The p-values are displayed in Fig 3 along with the raw spectra and grouped peak data for one of 33 significant features. When looking at the spectra that correspond to these features the difference between red and white wines is is obvious. However, manually searching the original spectra for these difference regions would be extremely tedious and time consuming. With speaq 2.0, this entire process takes about 3 minutes with 1 CPU and a mere 50 seconds with 4 CPUs (2.5 GHz machine).

### Onion intake in mice data

In the second validation we will compare the results of the presented method with those obtained by Winning et al. [29]. The objective of their study was to search for onion intake biomarkers in mice from 4 groups with increasing percentages of onion in their diet (0, 3, 7 and 10%). Subsequently urine was collected and analyzed with NMR spectroscopy. If the workflow can identify such onion intake biomarkers, it can possibly also be applied to search for other metabolic biomarkers.

### The struggle with bins and intervals

Winning et al. [29] used intervals methods (binning) to summarize the spectra. The internal workings of these interval methods are almost always the same: divide the spectra in intervals (a.k.a. regions, bins, buckets, etc.) and use these as variables. There are a number of problems with such methods, both at the pre-processing level (choice of the right boundaries, disappearance of relevant peaks because of noisy peaks in the interval) and at the results level (need for manually checking the relevant intervals for exact locations of interesting peaks).

### Towards a small and usable data matrix

Our method elegantly avoids these problems. The proposed method takes the raw NMR spectra (see supplementary files) and uses wavelets to convert these to peak data. The results are presented in Fig 4 (right half). Note that the authors of the study removed part of the data between 4.50 and 5.00 ppm because of insufficient water suppression [29] (see supplementary files). Next the peaks are grouped together. The grouped peaks are now called features. These features form the data matrix used as input for the statistical analysis. The dimensions of this data matrix are 31 samples by 677 features. This is a substantial reduction from the original spectra data matrix of dimensions 31 samples by 29001 measurement points.

**Fig 4.**
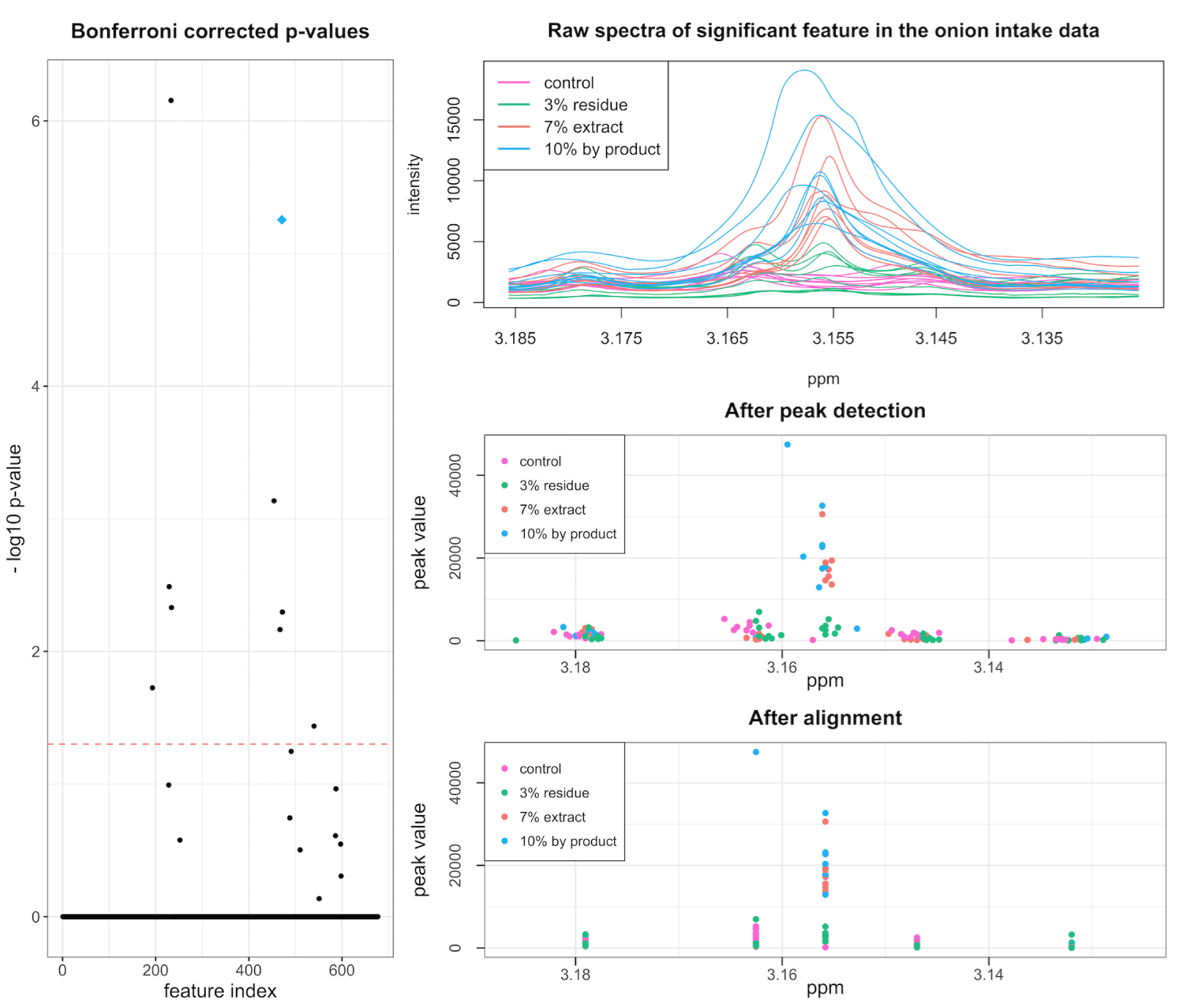
Differential analysis results of onion intake in mice data. (Left) the Bonferroni corrected p-values for the features resulting from the differential analysis and (right) one of the features with a significant p-value (indicated with the blue diamond on the left image): (top) raw spectra, (middle) data after peak detection and (bottom) data after grouping.

### No grouping is found by PCA

In line with the original analysis by Winning et al. the feature data matrix is Pareto scaled and centered. The results of the PCA analysis, as presented as a score plot in Fig 5, are analogous to those of [29]: there is no obvious clustering of the groups (not on onion intake class, nor on control vs onion intake).

**Fig 5.**
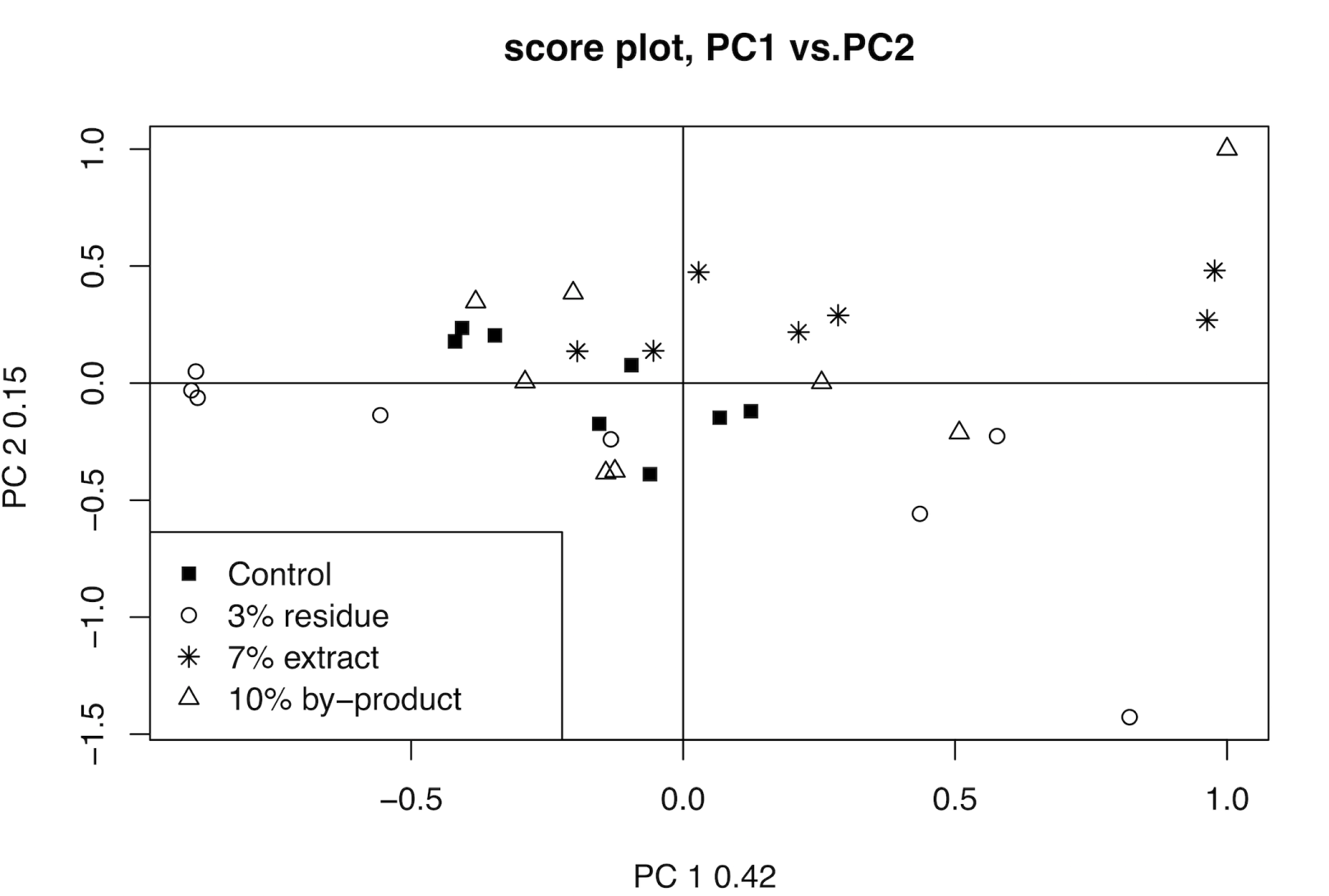
PCA analysis of onion mice data. The onion mice data matrix is Pareto scaled and centered. There are no clear trends present in the PCA results, this matches the results of Winning et al. [29].

### Locating possible biomarkers with ease

From this point onwards the merit of the wavelet based analysis becomes more obvious. Winning et al. have to resort to interval partial least squares (iPLS) and interval extended canonical variate analysis (iECVA), After careful cross validation these methods lead to intervals that have to be checked manually for interesting peaks. With our method the story becomes easier. The feature matrix is processed with the differential analysis method. In this case there are more than two groups. However this is not a problem for the differential analysis method since it is based on linear models and in this case there exists a numerical relationship between all the groups, (i.e. the percentage of onion in the diet). Every feature gets a Bonferroni corrected p-value assigned indicating how well the feature corresponds to the increasing onion concentration. The distribution of uncorrected p-values see the supplementary files. The corrected p-values are shown in Fig 4 along with an excerpt of one of the significant peaks. In total, 9 peaks were found to be significant. The 9 significant peaks can be used to search HMDB to find the possible biomarkers related to onion intake.

### Identifying the biomarkers

Merely submitting all peak ppm values to HMDB will not produce the correct outcome, as HMDB expects all peaks to correspond to a single metabolite. To avoid submitting peaks from multiple metabolites to an HMDB search, a correlation-based clustering step is performed on the highly significant peaks. The result from this clustering, based on peak intensity correlations, is displayed in Fig 6. The significant peaks are grouped into 5 clusters, where the minimal Pearson correlation between any two peaks in the same cluster is higher than 0.75. These peak clusters are used to search HMDB (tolerance ± 0.02), by submitting the ppm values of the peak groups within a cluster. When submitting the cluster of 4 peaks, the top hit is 3-hydroxyphenylacetic acid (HMDB00440) with a Jaccard index of 4/9. This molecule is also identified in the original paper as a biomarker for onion intake. However, in the original paper this is done only by looking at a small region around 6.8 ppm, as compared to the speaq 2.0 analysis which yields peaks in multiple ppm regions that can be used for identification. Note that in this case the correlation approach is not perfect, as the peak with index 18662 can actually also be assigned to 3-hydroxyphenylacetic acid (raising the Jaccard index to 5/9 upon also submitting this peak to HMDB). When the cluster that only contains peak 19723, with corresponding ppm value of 3.1558, is submitted to HMDB the top hits are dimethyl sulfone and 9-methyluric acid, both with a Jaccard index of 1/1. These results match those from the original paper where dimethyl sulfone (HMDB04983) is identified as a biomarker for onion intake. Raw spectra of the main peaks for both biomarkers are shown in the supplementary files.

**Fig 6.**
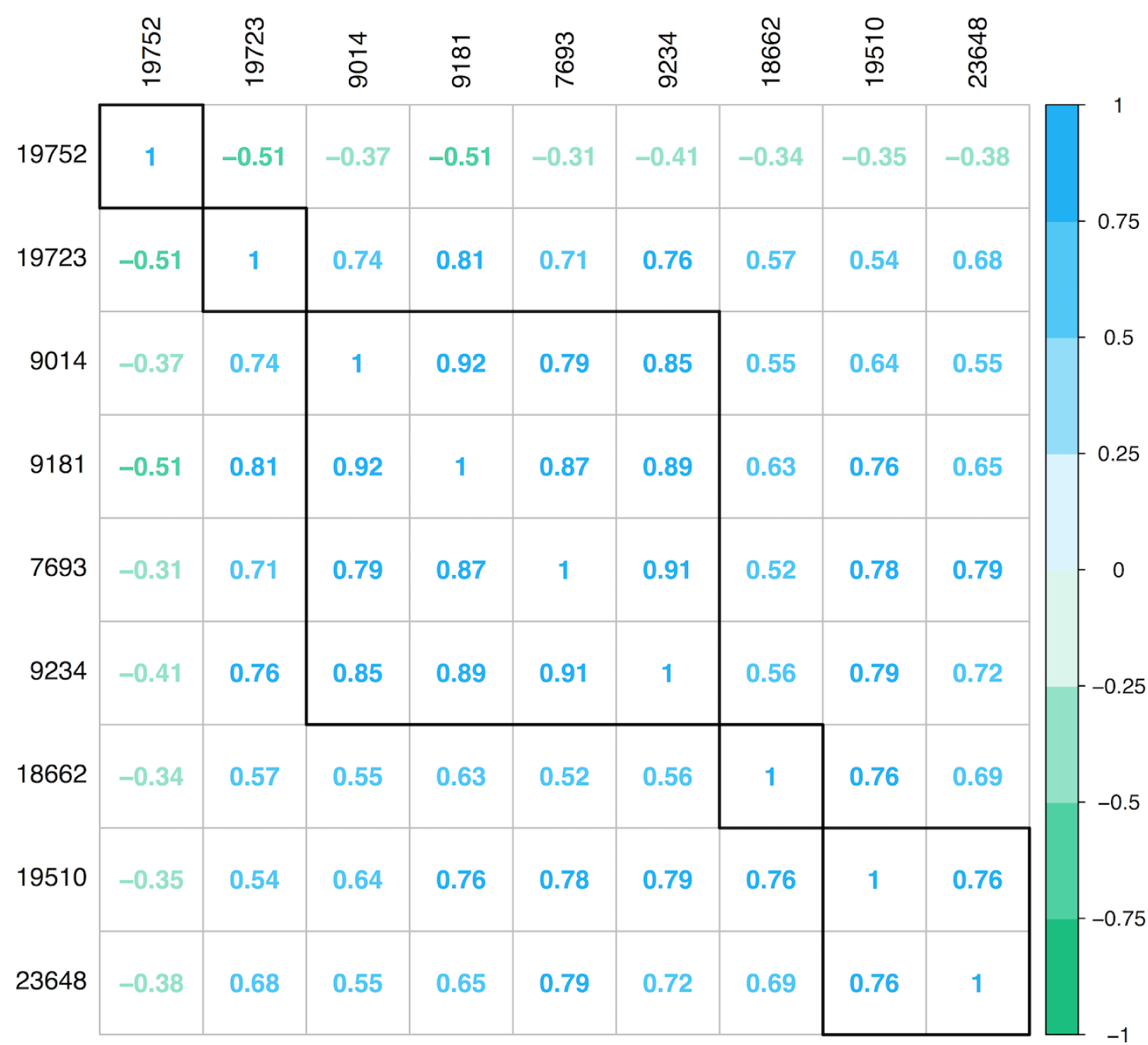
Correlation analysis of significant peaks. The significant peaks, which are indicated by their peakIndex value, are clustered based on their Pearson correlation. The group of four peaks correspond to the 3-hydroxyphenylacetic acid biomarker, peak nr 19723 corresponds to the dimethyl sulfone biomarker. Both biomarkers are also identified in the original analysis paper [29], but with only one peak for the first biomarker.

### The other peaks explained

The other peaks found cannot be identified querying HMDB. The peak with index 19510 is somewhat absorbed in the background. The peak with index 23648 ends up in a cluster with non-significant peaks that are assigned to ethanol within HMDB, when the correlation procedure is run on the entire dataset. As HMDB does not assign the 23648 peak to ethanol, this may indicate that this is a derivative or a byproduct of ethanol. The peak with index 19752 is actually a peak in the tail of the large peak of one of the identified biomarkers, namely dimethyl sulfone. The fact that this peak is significant is caused by an artifact of the wavelet based peak detection since it considers the tail of the large dimethyl sulfone peak as the baseline for the small peak. So when the dimethyl sulfone peak is larger, the baseline for the small peak is also larger and therefore the peak diminishes. This is also the reason why this peak is anti-correlated with the dimethyl sulfone peak.

## Discussion

We present an easy way of converting 1D NMR spectra (or other 1D spectra) to peak data by using wavelets for peak detection. This wavelet based method performs better than binning or other spectra summarizing methods as the dimension of the dataset is greatly reduced with little to no loss of information, while requiring no user intervention. After the wavelet based step the peaks are grouped via a hierarchical clustering method. These groups of peaks are called features. The features can easily be analyzed with a myriad of statistical techniques or data mining approaches. Our method has been implemented in an entirely new version of the existing speaq *R* package, in order to provide an entire solution for easy 1D NMR data analysis. As each step in the workflow is available as a single function, analysis pipelines can be constructed easily and with little additional user interaction, fostering improved research reproducibility and shareability

Besides the possibility to perform a complete standalone analysis, our method can also be used in tandem with other commonly used tools that rely on summarized spectra. Specifically, it can be used to quickly and efficiently produce a high quality peak list. Such a peak list is the starting point of an analysis with for example the often used MetaboAnalyst [14].

The data processed in this article came in a matrix format with ppm values and intensities. Other proprietary software or open-source frameworks are thus needed if only the raw NMR Free Induction Decay signal (FID) is available and conversion to the frequency space is needed. Optional pre-processing steps on the raw FID signal like zero-filling, apodization, and phase-shifting have to be performed prior to employing speaq 2.0, if they are desired. These pre-processing steps along with compatibility with the open source nmrML (http://www.nmrml.org) format are on the road-map for future developments.

We expect the introduced method to be especially useful for processing NMR spectra from large cross-platform experiments that combine NMR and LC-MS. Often software packages like XCMS [5] are used to process LC-MS data. These open source packages also employ the standard paradigm of peak-picking, grouping, etc. so the integration of data or results should be facilitated with this framework. The method in itself also has merit as is clearly demonstrated in the case of the onion intake in mice data. The analysis is fast, sensitive to both small and large peaks and user-independent. Also, when comparing the results we obtained to the work presented by Winning et al. [29], our analysis required less user interaction and yields more peaks in the end that can be used to identify the possible biomarkers, resulting in an improved confidence in the results.

The user-friendliness of speaq 2.0 should also allow people with little experience in R to use the package. To this end, the code for both the performed analysis has been made available on CRAN (https://cran.r-project.org/package=speaq) and GitHub (https://github.com/beirnaert/speaq) as a starting point. Also, it can serve as an attractive option for researchers interested in switching from closed, proprietary software to open-source, especially if the goal is to speed up analysis, improve reproducibility and increase control over workflows and algorithms.

## Acknowledgments

This research is supported by a GOA (Geconcentreerde Onderzoeksactie) from the University of Antwerp. Additional thanks go out to Aida Ligata (Mrzic) for the many fruitful discussion on figures and visualizations. Also we would like to thank all users of the speaq R package for their valuable feedback and support.

